# High temperature molecular motions within a model protomembrane architecture

**DOI:** 10.1101/2022.03.31.486527

**Authors:** Loreto Misuraca, Tatsuhito Matsuo, Aline Cisse, Josephine LoRicco, Antonio Caliò, Jean-Marc Zanotti, Bruno Demé, Philippe Oger, Judith Peters

**Affiliations:** Univ. Grenoble Alpes, CNRS, LIPhy, 38000 Grenoble, France; Institut Laue Langevin, F-38042 Grenoble Cedex 9, France; Institute for Quantum Life Science, National Institutes for Quantum Science and Technology, 2-4 Shirakata, Tokai, Ibaraki, 319-1106, Japan; INSA Lyon, Université de Lyon, CNRS, UMR5240, Villeurbanne, France; Institut Universitaire de France; Laboratoire Léon Brillouin, CEA, CNRS, Université Paris-Saclay, CEA Saclay, 91191 Gif-sur-Yvette Cedex, France

## Abstract

Modern phospholipid membranes are known to be in a functional, physiological state, corresponding to the liquid crystalline phase, only under very precise external conditions. The phase is characterised by specific lipid motions, which seem mandatory to permit sufficient flexibility and stability for the membrane. It can be assumed that similar principles hold for proto-membranes at the origin of life although they were likely composed of simpler, single chain fatty acids and alcohols. In the present study we investigated molecular motions of four types of model membranes to shed light on the variations of dynamics and structure as a function of temperature as protocells might have existed close to hot vents. We find a clear hierarchy among the flexibilities of the samples, where some structural parameters seem to depend on the lipids used while others do not.

## Introduction

The onset of first cellular life on Earth goes back to about 4 billion years ^1^. Therefore, the origin and nature of the first cell membranes and their constituents are not fully determined. Because of the supposed lack of molecular complexity on the early planet, the first cell membranes were most likely composed of simple, single chain fatty acids ^2^ and fatty alcohols, raising questions as how they could withstand the very variable and extreme surrounding environment ^3^ of the oceanic hydrothermal vents or fields. One of the commonly accepted scenarios for the origin of life is indeed related to such places ^4^. In particular, the membranes may have had to cope with high temperatures (typically around 85°C) and high hydrostatic pressures (up to a few hundred bars), and simultaneously accomplish elementary but crucial functions of cell membranes: divide the inside and outside into separate compartments, and assure a limited permeability combined with a certain stability and flexibility.

As proto-cells or proto-membranes are no longer accessible due to evolution, the only way to probe the properties of these hypothetical systems is to re-construct membranes from simple lipids with short chain lengths, presumably favoured by prebiotic synthesis ^5-7^, and to expose them to the supposed environmental conditions. Here we concentrate on fatty acids and fatty alcohols and phospholipids, all with chain lengths of 10 carbons (named hereafter C10), which are among the shortest to self-assemble into lipid bilayers ^8, 9^.

Specifically, we study here several possibilities (see Figure 1): first, decanoic (or capric) acid, which is a C10 fatty acid known to form vesicles under fixed conditions ^9^. Second, a possible architecture for protocell membranes, consisting of a 1:1 mixture of capric acid with a fatty alcohol of equal chain length (decanol). These short, single chain amphiphilic molecules were in fact the ones readily available in the prebiotic environment and have been found able to form stable vesicles ^10^. In a recent work, we reported the remarkable properties of this membrane architecture which allow it to withstand extreme temperature conditions ^11^. Indeed, such a combination of fatty acids and alcohols improves the vesicles’ stability at room temperature and it induces a high-temperature conformational change at T ≥ 60 °C that leads to vesicle fusion with stable membranes at temperatures as high as 80 °C. In the following, this model membrane will be referred to as C10mix.

**Figure 1:**
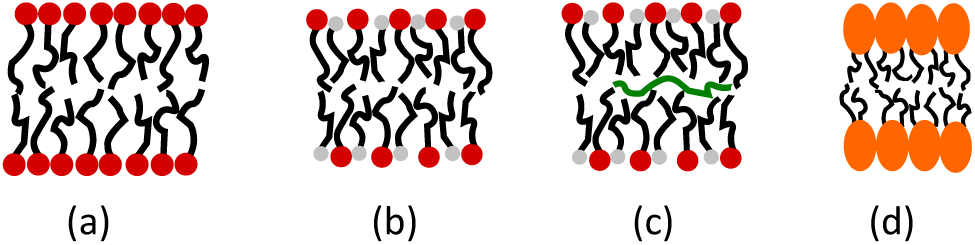
Four models for protomembranes: (a) Capric acid, (b) Capric acid + decanol (C10mix), (c) C10mix + eicosane, (d) DCPC.

The third model is inspired by the work of Cario et al. ^12^. These authors proposed an important role of long chain isoprenoid alkanes (referred as “apolar lipid”) in the membrane mid-plane to further trigger its physico-chemical properties in relation to extreme conditions. In fact, Salvador-Castell et al. did find that they had a stabilizing role in model archaeal membranes ^13^ and Misuraca et al. reported an increased stability of lipid membranes in presence of alkanes ^14, 15^. Thus the C10mix was enriched with small percentages of eicosane, a linear alkane with 20 carbons (C20). The C10mix containing eicosane can be considered as the most complete prebiotic model that sums up the requested features of molecular abundance and need for a strategy against high temperature.

To further compare the decanoic acid to a more evolved lipid, a phospholipid having two C10 acyl chains, but the much larger choline headgroup was chosen: 1,2-didecanoyl-*sn*-glycero-3-phosphocholine (DCPC). The acyl chains are bound to the headgroup through a glycerol backbone, which can stiffen the structure.

Not only the structural composition of lipidic membranes is significant for their behavior, but their dynamic properties have an equally important impact ^16^. High temperature is known to increase the thermal energy and thus molecular motions within membranes, hence jeopardizing the stability of the system. Nonetheless, to be functional a cell membrane requires a specific state of fluidity and therefore high flexibility. Both together can be ensured by molecular dynamics within the membranes.

Many kind of motions were identified within lipidic membranes, from small local motions up to collective motions of the whole system. A well adapted technique for their observation and analysis is incoherent neutron scattering, which probes essentially the motions of hydrogen atoms and of the molecular subgroups to which they are bound ^17-20^. Neutron instruments give access to a broad range of energy resolutions, which are inversely propotional to the accessible time window, and thus allow different types of motions to be probed. In the present study, we combine elastic incoherent neutron scattering (EINS) experiments giving access to atomic mean square displacements (MSD) with quasi-elastic neutron scattering (QENS) experiments to characterize the kind of motion present in the sample and the associated amplitudes. To the best of our knowledge, such complete investigation on multilamellar vesicles of one type of lipids, of mixed lipids and alcohol or in presence of alkanes was never attempted before, mainly due to the lack of a suitable model to describe the dynamics. However, as molecular dynamics are so important for the understanding of the correct functioning in a certain state, we show here which parameters can be extracted, which ones are the most impacted by the structural differences and how we can interpret these variations.

## Experimental section

### Sample preparation

Decanoic acid, 1-decanol, perdeuterated eicosane, DCPC, bicine buffer and D_2_O were purchased from Sigma Aldrich (Merck). The samples were prepared by dissolving the decanoic acid and the decanol in a CHCl_3_ solution and DCPC in chloroform:methanol (2:1), followed by drying under a stream of nitrogen. The samples were then placed in a desiccator and left under vacuum overnight. The sample weights were checked at each step, to make sure that all the organic solvent was evaporated. For the samples consisting of decanoic acid : 1-decanol (C10mix) the final ratio was 1:1.

The bicine buffer was prepared at a concentration of 0.2M by dissolving hydrogenated bicine in D_2_O, following previous protocols ^10^. The buffer was filtered with a 0.2 μm millipore membrane before use. The dried organic solutions were then resuspended in buffer by vigorous vortexing for ≈ 1min, leading to the final milky-foamy solutions characteristic of the presence of large multi-lamellar vesicles (MLVs). The samples to be measured were then diluted in D_2_O buffer to a final concentration of 60 mg/ml (350 mM). The samples were placed in flat aluminium sample holders and sealed with indium. Their weights were checked before and after the experiment to control that no solution was lost.

### Incoherent neutron scattering

The incoherent neutron scattering cross section of hydrogen exceeds by far that of all other atoms present in biological systems ^21^ and also that of its isotope deuterium. Therefore, contrast variation can be used to highlight specific parts of the samples, as in our case the lipids rather than the buffer or the alkanes, whose presence will only be accounted for by their impact on the membrane dynamics.

For our study, we used the spectrometer IN6-Sharp operated by the Laboratoire Léon Brillouin (Saclay, France) at the Institut Laue Langevin (ILL) in Grenoble, France. IN6-Sharp is a cold neutron time-of-flight ^22^ with an energy resolution of 70 μeV at 5.1 Å incident wavelength, giving access to a time window of about 10 ps. Elastic scans were performed from 278 to 355 K with a temperature step of 1 K and a measuring time of 2 minutes per point. QENS measurements were conducted on the same samples at the temperature points of 278, 293, 323 and 353 K for 2.5 hours each. Additionally, the bicine buffer was measured in the same conditions with the exception that the QENS scans were acquired for only 2 hours due to time limitations. An empty cell and the completely incoherent scatterer vanadium were measured at room temperature for correction and normalisation purposes. The empty cell and buffer data were subtracted from the sample data taking into account their different absorptions based on the correction formula of Paalman−Pings coefficients ^23^. Vanadium data were used to normalise the elastic data and to determine the instrumental resolution for QENS data. Data reduction for EINS was carried out using the LAMP software ^24^ available at ILL and using the IGOR Pro software (WaveMetrics, Lake Oswego, OR, USA) for QENS.

### Elastic incoherent neutron scattering

EINS measurements give access to the elastic incoherent structure function *S*(*Q*, 0 ± *ΔE*), where *Q* is the modulus of the difference between the incoming and outgoing neutron wavevectors and *ΔE* the energy resolution of the instrument. The structure factor can be written, using the Gaussian approximation ^25^, which assumes that the distribution of the atoms around their average position follows a Gaussian distribution,

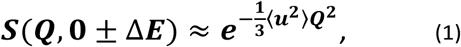

where the *<u*^*2*^*>* are the MSD. As *Q* approaches zero, the approximation is strictly valid and it holds up to ⟨*u*^*2*^⟩*Q*^*2*^ ≈ 1 ^26^.

The average MSD can be obtained from the slope of the logarithm of the scattered intensities according to

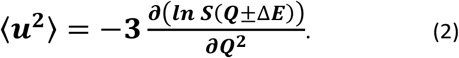

The *Q*-range used for fitting was on IN6-Sharp 0.45 to 1.4 Å^−1^. It is possible to further determine the force constant or resilience <*k*> from the slope of the MSD ^27^ through

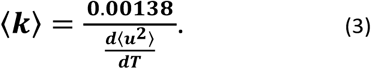

Here <*k*> is expressed in Newton per meter when <*u*^*2*^> is given in Å squared and T is the absolute temperature.

### Quasi-elastic neutron scattering

Whereas elastic scattering permits only averaged local motions to be seen, QENS is the tool to separate different kind of movements and to characterize them. For that, each spectrum can be fitted after the data reduction with a phenomenological equation containing terms which account for various diffusions:

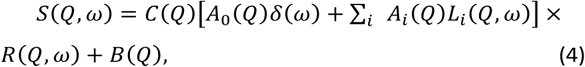

where *ω* is the energy transfer in multiples of ħ, *C(Q)* is a scaling factor to adjust the model and the absolute data, and *A*_*0*_*(Q) δ(ω)* is the *Q*-dependent elastic incoherent structure factor (EISF). The functions *L*_*i*_*(Q,ω)* are Lorentzian functions of half-width at half-maximum (HWHM*) Г*_*I*_, each describing a certain type of diffusive motion in the lipids. Their amplitudes *A*_*i*_*(Q)* are also *Q*-dependent. *R(Q,ω)* refers to the instrumental resolution which is obtained from the vanadium measurements. *B(Q)* is a flat background. The sum can theoretically be extended over an infinity of contributions, but real data allow only to resolve a few of them. Careful checks determined that 3 Lorentzians were sufficient to reasonably fit the data here without overfitting them. They represent slow (e.g. two-dimensional diffusion within the membrane plane and one-dimensional diffusion of the whole lipid perpendicular to the plane), intermediate (e.g. rotational diffusion of the whole lipid, flip-flop and rotational motions of the headgroup) and fast motions (e.g. two-dimensional diffusion of the tail group and jump-diffusion of hydrogen atoms in methyl groups), as classified in the Matryoshka model recently proposed for a very complete description of motions within a membrane ^28^.

The behaviour of *Г* as function of *Q*^*2*^ is indicative for the type of movement ^29^: a linear behaviour represents translational diffusion characterized by the diffusion coefficient *D*_*T*_:

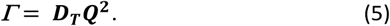

Sometimes, *Г* is not crossing the origin for Q → 0, what is the signature of confinement for larger amplitudes. In this case, it is advised to fit the data only from a Q_min_ on, where the linear behaviour sets in.

A constant behaviour is typical for rotational diffusion characterized by the rotational diffusion coefficient D_R_:

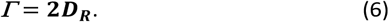

Finally, jump-diffusion is typical for infinitely small, elementary jumps and a certain time spent by the protons in between, the residence time *τ*, in addition to the jump diffusion coefficient D_jump_. Here, *Г* as function of *Q*^*2*^ increases towards a constant limit, *Г*_*∞*_ *= 1/τ*:

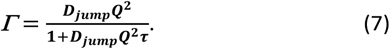

We fitted the experimental spectra in the *Q*-range from 0.37 to 2.02 Å^-1^ and within energy transfers ω from -10 meV to 2 meV (see an example in Figure 2).

**Figure 2:**
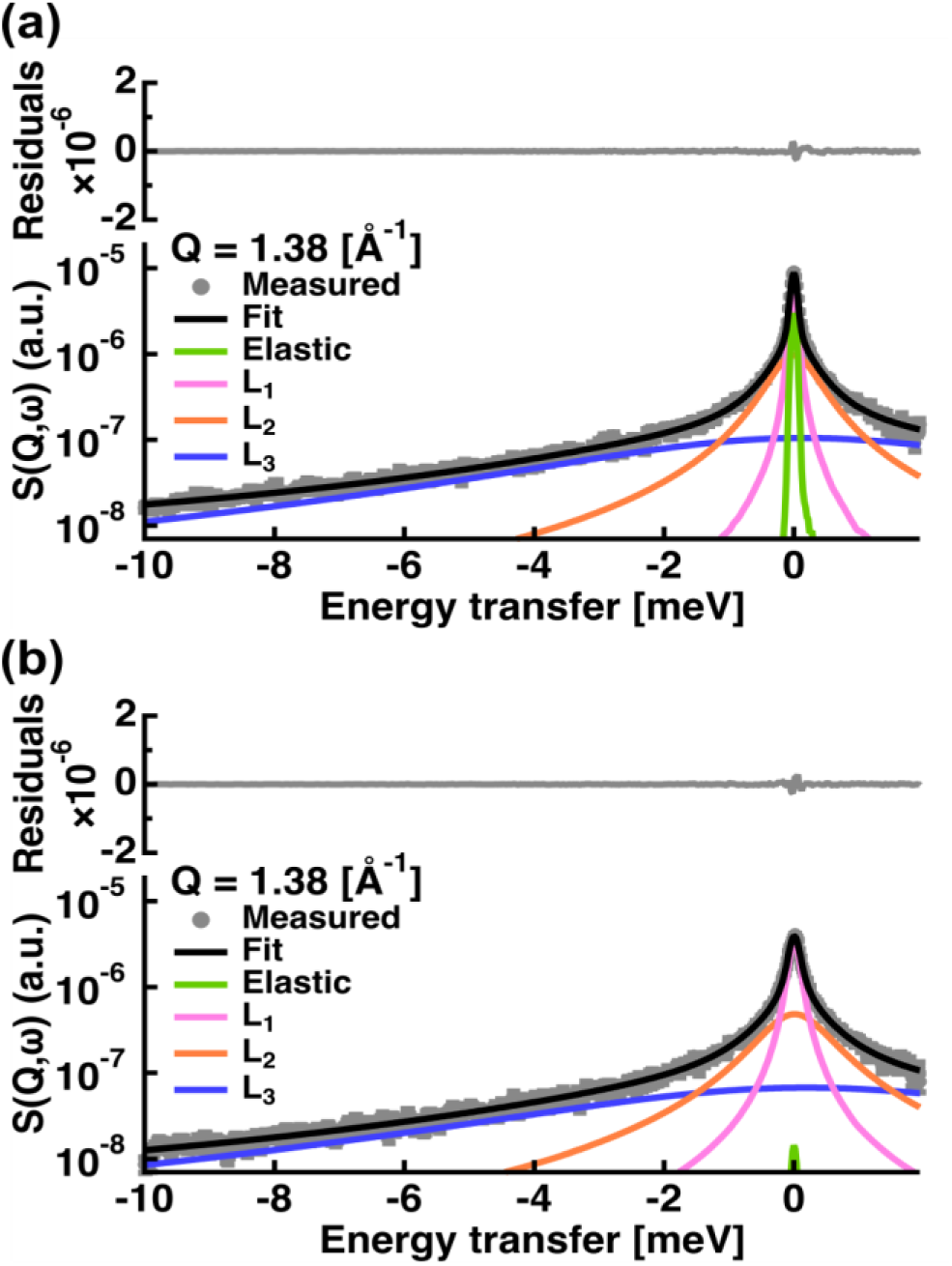
Examples of fit results for the spectra of (a) capric acid and (b) C10mix, both at *Q* = 1.38 Å^-1^ and T = 293 K. The grey circles are the data points with errors, the black line is the total fit line. The green line represents the elastic contribution convoluted by the resolution function. The magenta, orange and blue curves are the Lorentzian functions corresponding to slow, intermediate and fast motions, respectively, convoluted by the resolution.

## Results and discussion

Experiments were performed in August 2020 on IN6-Sharp (DOI: 10.5291/ILL-DATA.CRG-2728).

### EINS

The integration of the elastic peak allowed the extraction of the MSD as a function of temperature through eq. (2) (see Figure 3). Generally speaking, lipids are very flexible molecules and our results are in close agreement with other examples in the literature (see reference ^30^, but note that in this reference the use of a different factor for the MSD).

**Figure 3:**
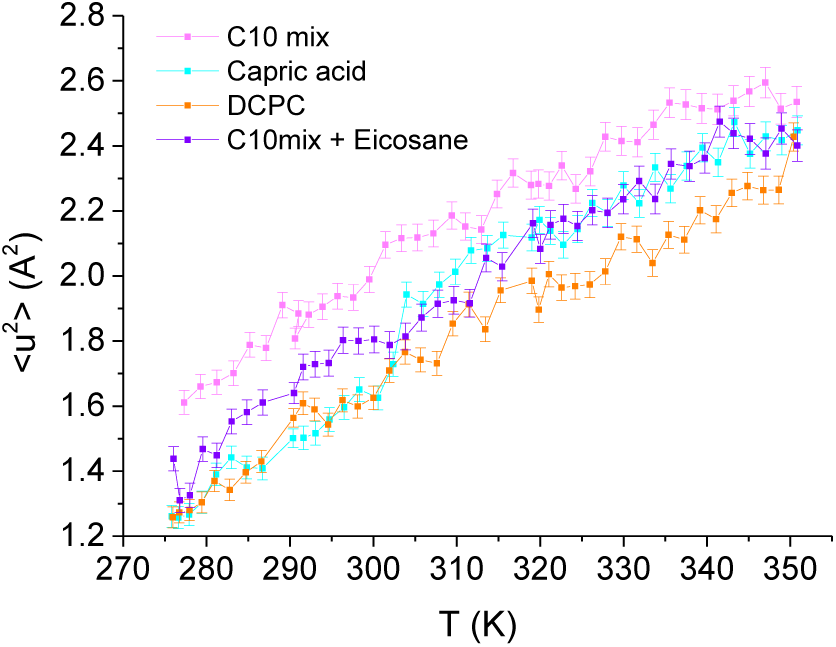
MSD for MLVs of the four samples as a function of temperature.

The slopes (<*k*>) of all four curves present very similar trends (see table 1), indicating that the four systems have similar stabilities. On the contrary, the MSD data highlights a few particularities for each one. As can be seen in Figure 3, the C10mix is the most flexible sample over the whole temperature range. However, we do not observe an impact of the conformational transition of the C10mix above 333 K on the local dynamics as seen by DLS ^11^. By contrast, the capric acid seems to undergo a steep raise in MSD at around 305 K, indicative of a phase transition, which is somewhat higher than the one observed by DSC in previous studies (287K) ^11, 14^. We interpreted the phase at lower temperature as a collapsed phase, which melts around this temperature. The shift in temperature observed here can have various reasons such as different temperature ramps for the two techniques or differences in hydration ^30^.

**Table 1:** The force constants of the four samples.

| | $\langle k \rangle$ (N/m) | $\Delta \langle k \rangle$ (N/m) |
| --- | --- | --- |
| C10mix | 0.102 | 0.003 |
| C10mix + eicosane | 0.092 | 0.002 |
| DCPC | 0.096 | 0.002 |
| Capric acid (< 305 K) | 0.085 | 0.005 |
| Capric acid (>305 K) | 0.124 | 0.007 |

In presence of eicosane, the C10mix becomes less flexible, as expected due to the hindrance of motions by an obstacle in the midplane of the bilayer, but as mentioned above, it maintains a stability similar to the C10mix as indicated by the slope values. Finally, DCPC presents the lowest MSD, only coinciding with those of capric acid at the lowest temperatures. The bigger headgroups, and the connection of the two C10 chains to the glycerol backbone could indeed hinder motions at the level of the heads or within the hydrophobic region.

### QENS

The treatment of the QENS data gave access to different information: First, we extracted the amplitudes *A*_*0*_ to *A*_*3*_ of all samples (see Figures S1 to S4 in the Electronic Supplementary information (ESI)). The amplitude *A*_*0*_ or EISF represents, at the highest *Q* value available, the proportion of particles seen as immobile within the instrumental resolution. At the two highest temperatures, 323 and 353 K, all four curves converge quickly to zero at higher Q values, indicating the high mobility. At the lower temperatures, capric acid and DCPC present values between 0.05 and 0.2 showing that not all hydrogens are participating in the motions yet. Capric acid has the highest values for these temperatures, but joins the other curves in between 293 and 323 K, in accordance with the phase transition found from EINS. The C10mix seems again the most flexible, but very similar to its counterpart with eicosane. The amplitudes *A*_*0*_ to *A*_*3*_ will be treated in more detail with the Matryoshka model ^28^ to extract also structural information (see below).

Second, the HWHM *Г*_*I*_ are representative of three identified types of motions (see Figures S5 to S7 in the ESI). It can be deduced by the different energy ranges they cover: approximately from 0.05 to 0.35 meV for *Г*_*1*_, from 0.25 to 1.25 meV for *Г*_*2*_ and from 3 to 8 meV for *Г*_*3*_. From their behaviours as a function of *Q*^*2*^, one identifies that *Г*_*1*_ corresponds to continuous diffusional motions, including confinement for C10mix and C10mix + eicosane, *Г*_*2*_ to jump diffusion and *Г*_*3*_ to rotational diffusion. The values of *Г*_*2*_ also fluctuate sensibly below the *Q*-value of 1.1 Å^-1^ around an almost constant value. Such attitude is indicative of a confined motion, which can be characterized by a radius of confinement *a*, to be determined through *a = π/Q*_*0*_, where *Q*_*0*_ stands for the momentum transfer where jump diffusion sets in. In the present cases, this critical value of *Q*_*0*_ was the same for all samples and equal to 2.9 Å.

With respect to the global translational diffusion coefficient of the lipids within the membranes (see eq. (5)), as shown in Figure 4 as a function of temperature for the four samples, it seems to be very similar in almost all cases except for the sample C10mix with eicosane, which presents a much higher value at high temperature. Moreover, the *Г*_*1*_ of this sample at 353 K shows a behaviour resembling more jump diffusion (increasing towards a limit and a constant value at the low-Q limit) than pure translational diffusion (linear behaviour), likely because of the presence of the alkanes which induces the confinement of the acyl chains, and limits the amplitudes of the motions. All values of the diffusion coefficients are in between 0.3 10^−5^ cm^2^/s at 278K and 2 10^−5^ cm^2^/s at 353K and lie below the value of bulk water, 2.3 10^−5^ cm^2^/s at 298 K and 6.81 10^−5^ cm^2^/s at 353K ^31^, which represents somehow the upper limit for such diffusional processes. site jump diffusion could be more appropriate here. The hierarchy of the residence times is consistent with the EINS results, e.g. C10mix < capric acid < C10mix + eicosane < DCPC. This quantity, representative for interactions between molecules, decreases with temperature as the lipids become more and more mobile and is highest for DCPC.

**Figure 4:**
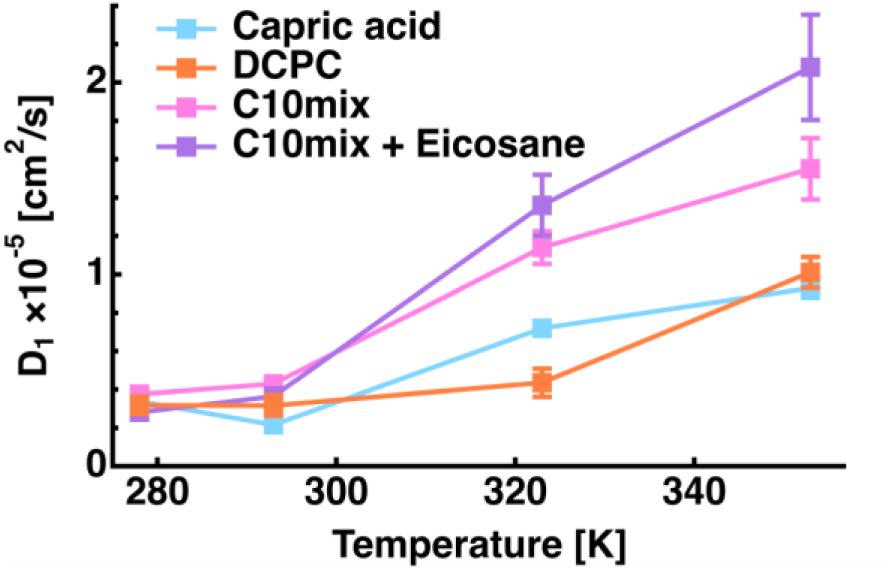
Global translational diffusion coefficient of all four samples as a function of temperature.

The last *Г*_*3*_ permits to calculate the rotational correlation time τ_3_, defined as the inverse of Г_3_, which is represented in figure S8 of the ESI. All values are very close and almost independent of temperature. These motions are the fastest with smallest amplitudes and thus almost not prone to the specific environment.

We further used a recently developed model to interpret the amplitudes *A*_*0*_ to *A*_*3*_ of eq. (4), the Matryoshka model ^28^ (see Figure 6). Although the Matryoshka model is dynamic, fitting the data with it, starting from values found in the literature, it allows to determine the impact of molecular dynamics on certain structural parameters. The amplitudes are globally fitted for each sample, but independently for each temperature point.

**Figure 5:**
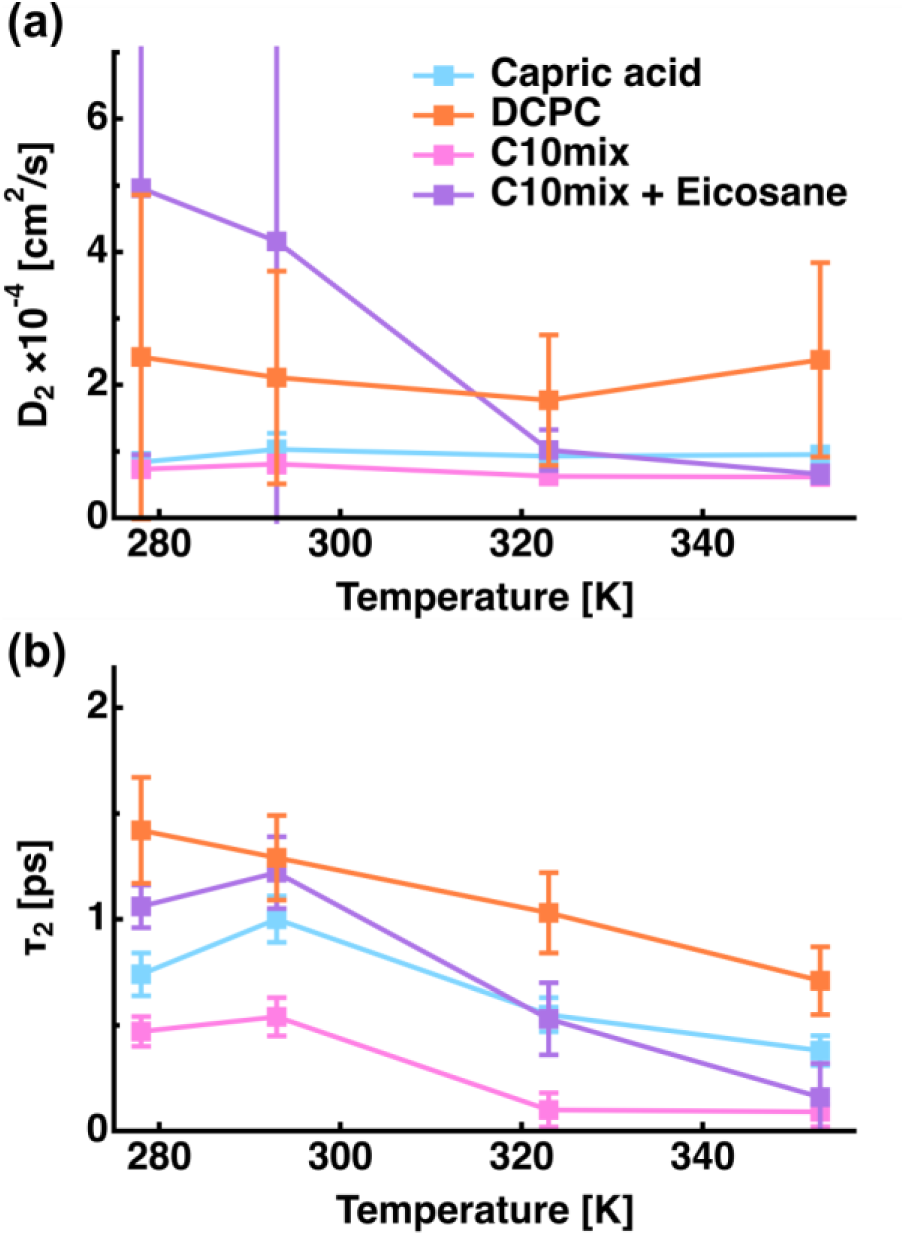
The jump diffusion coefficient and the residence time for all samples as a function of temperature extracted from *Г*_*2*_.

**Figure 6:**
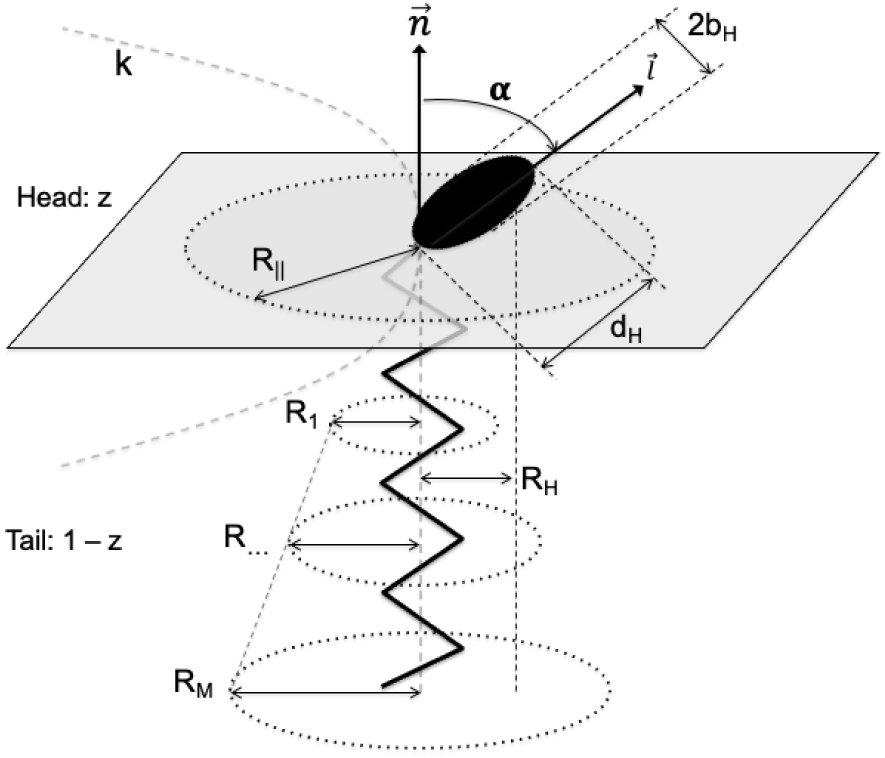
Representation of a lipid molecule within the Matryoshka model and fit parameters.

The second type of motion corresponds to jump diffusion and allows the determination of the jump diffusion coefficient and residence time according to eq. (7). Figure 5 represents these quantities for all samples. However, this model does not describe very well the motions within C10mix + eicosane and DCPC, leading to huge error bars. We can speculate that 2, 3-

The Matryoshka model describes the two alkyl chains as one and the same cylinder. The modelled parameters are the following: *R* is the lateral diffusion radius of the whole lipid in the membrane plane. *k*_*force*_ represents a force constant or resilience characterising the relaxation of the lipid with respect to out-of-plane motions. The whole lipid can rotate, expressed through the distance head-lipid *R*_*H*_. Flip-flop motions of the head group are described by the angle α (we are using the definition of Pfeiffer et al. ^17^ and not the exchanges of lipids from one layer to the other, which correspond to much longer time scales) and the heads with radius *b*_*H*_ can rotate around their own axis. *R*_*1*_ stands for the minimum radius of diffusion of the lipid tail, which according to the distance from the head increases with 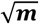, *m* being the index of the methylene group position on the tail and *M* the total number of these groups (here: *M* = 10 for all samples). H atoms inside the methylene and methyl groups can undergo jump-diffusion. The parameters needed to describe these motions are the distance H-C-H *d* between the two-sites and the probability of jump events *ϕ*. The model depends further on experimental parameters known for lipids as the number *z*, which is the proportion of H atoms in the head (here *z* = 0.05 for capric acid, = 0.03 for C10mix with and without the eicosane, = 0.32 for DCPC) and on the fraction of H atoms, seen as immobile within the instrumental resolution, termed here *p*_*imm*_. For a validation of the model upon real data, see _32, 33_.

In figure S9 of the ESI we show the fits of the amplitudes for all four temperatures of our samples. Globally, they work remarkably well, giving mainly rise to reduced *χ*^*2*^ values between 1 and 16. This is more satisfying as the model was first developed for phospholipids and successfully applied here for a first time to single chain fatty acids and alcohols, too.

The extracted fit parameters are presented in Figure 7. Some remarks are noteworthy: When *z* has a very small value (as here for capric acid and C10mix), it is almost impossible to model the head motions, which explains the huge error bars for parameters *b* and *α* for these samples. For DCPC, *α* was fixed to 30°, similarly to DMPC, as the head groups are the same ^32^. In that case, the error bars are smaller and *b* is in the same order of magnitude as for DMPC ^32^, which is consistent with identical head groups. The other quantity for which the fits cannot give reliable results is the tilt angle *α*. In fact, the head group of capric acid and C10mix is so small that its orientation does not make sense.

**Figure 7:**
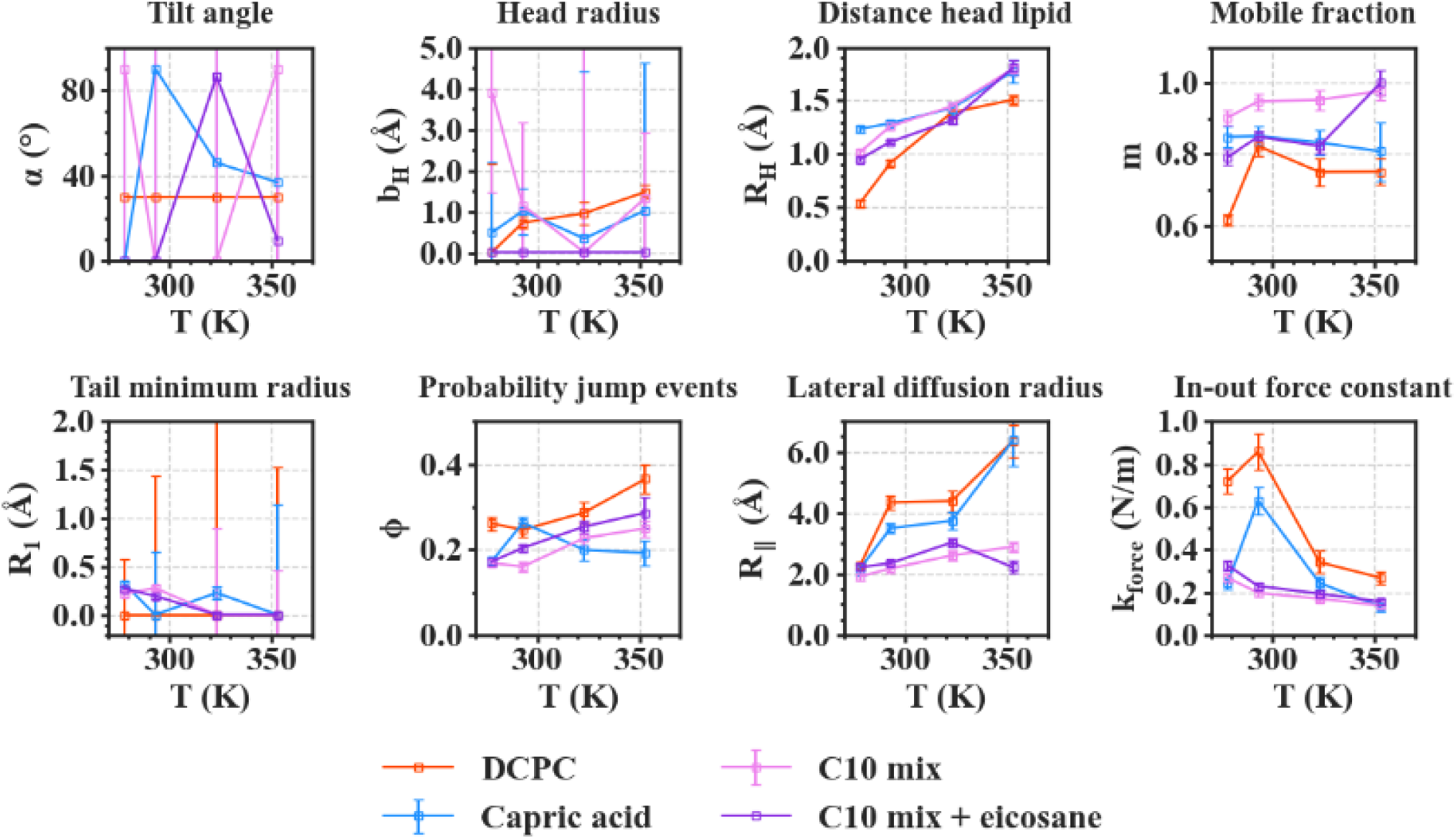
Fit parameters obtained for all samples as a function of temperature. Error bars are within symbols if not shown.

In contrast to DMPC ^27^, *R*_*1*_ is close to 0 for almost every sample, meaning that the corresponding motion, the 2D diffusion of the effective tail, is negligible compared to the jump-diffusion of methyl groups. As it occurs for all systems with different head groups, this is probably related to the smaller chain length (*M* = 10, compared to *M* = 14 for DMPC), which would hinder the 2D diffusion of the tail. However, *R*_*1*_ seems slightly bigger for DCPC, which can be assigned to the lower phase transition temperature of DCPC so that these lipids are in a liquid crystalline state even at 293 K.

The jump-diffusion, represented by the *ϕ* parameter, seems to be more likely to occur in DCPC, and much less in capric acid. This could be due to the larger head group and the liquid crystalline state of DCPC, allowing more motions within the chains, or to the fact that there are two tails in DCPC, compared to one in all other systems, and one effective tail considered in the Matryoshka model (as discussed by Bicout et al. ^16^). Eicosane does not seem to have a strong effect on this parameter, which is reasonable as it is supposed to be mainly in the midplane of the bilayers, and thus is not supposed to interact with the majority of methylene and methyl groups.

Overall, the fraction of mobile H atoms is similar for all samples, and is about 80 %. Notably, the hierarchy between the samples is close to the one encountered for the MSD in Figure 3, where the C10mix displays the highest mobile fraction, and DCPC the lowest. We also retrieve the lowering effect of eicosane, with a decrease in the mobile fraction, except for the highest temperature.

Concerning the motions of the whole lipid, we can first comment on the rotation, corresponding to the parameter *R*_*H*_. While increasing with temperature, following the increase in thermal energy, the values of *R*_*H*_ stand for all samples around 1.5 Å, which can be put in comparison with measurements by SANS and NMR. Indeed, they give an estimated value for the lipid height of 1.7 Å ^11^. Here *R*_*H*_ is close to the projection of the lipid height, which is then supposed to be slightly smaller. It is slightly smaller for DCPC than for the other samples, because the single chain amphiphiles will rotate along their longitudinal axis much more easily than DCPC ^34^.

For the lateral diffusion radius and the in-out force constant, one recognises two groups, formed by DCPC and capric acid on one side and C10mix with or without eicosane on the other side. We can conclude that in the latter case, in-plane motions of the lipids are more restricted, but the membrane is more flexible in the out-of-plane direction. The presence of eicosane can certainly explain such behaviour, but it seems already present in the C10mix alone. One might speculate that it favours the requirements of a protomembrane.

## Conclusions

In summary, the exact composition of the lipid model membrane has a clear effect on molecular dynamics and structural parameters. EINS established the existence of a phase transition around 305 K for capric acid, likely corresponding to the melting of a collapsed phase already seen earlier ^11^. Otherwise we found the following hierarchy in the flexibilities with C10mix > capric acid > C10mix + eicosane > DCPC, although the stabilities (determined by the force constants, table 1) were very similar for all of them. It fosters the idea that the C10mix permits a high degree of adaptation to the environment due to the flexibility, which is however slightly decreased by the presence of the alkanes located in the midplane. As shown earlier ^14, 15^, the alkanes on the contrary increase the membrane’s stability under extreme temperature and pressure conditions.

QENS measurements revealed that the C10mix sample containing eicosane presented a very high translational diffusion at the highest temperature (which is supposed to correspond to the native condition). In contrast to that, the jump diffusional coefficient, characteristic of smaller localised motions, of this sample evolved from a high value at low temperature to the same value as for capric acid and C10mix alone at high temperature. Thus such a small value seems mandatory for a good functioning. All these values are quite different from those of DCPC, which is a non-prebiotic phospholipid with a much bigger headgroup.

The amplitudes of the motions permitted further structural parameters to be obtained through a fitting procedure with some starting values taken from the literature, as observed through the dynamical changes. The two parameters *b* and *α* are not well defined for capric acid and C10mix due to their small head groups and are fixed for DCPC. Some parameters, such as *R*_*H*_, *R*_*1*_ and *φ* do not differ significantly among the samples. On the contrary, *m, R*_*‖*_ and *k*_*force*_ show respectively two groups of samples, assembling capric acid and DCPC on one side and C10mix and C10mix with eicosane on the other side. So one might speculate that these three parameters are the most important to guarantee the good functioning of a membrane built from short-chain lipids and alcohol, with or without alkanes in the midplane. We retain a higher mobile fraction for these samples, especially at higher temperature, paired with a smaller lateral diffusion radius and a lower in-out force constant, indicating the motional direction perpendicular to the surface being favoured over lateral motions.

## Acknowledgements

This work was funded by the French National Research Agency program ANR 17-CE11-0012-01 to P.O. and J.P. L.M. was supported by a scholarship from the Institut Laue-Langevin (ILL) PhD program. The authors thank ILL for neutron beamtime on IN6-Sharp (DOI: 10.5291/ILL-DATA.CRG-2728).

