## Supplementary Information for "High temperature molecular motions within a model protomembrane architecture"

Electronic supplementary information

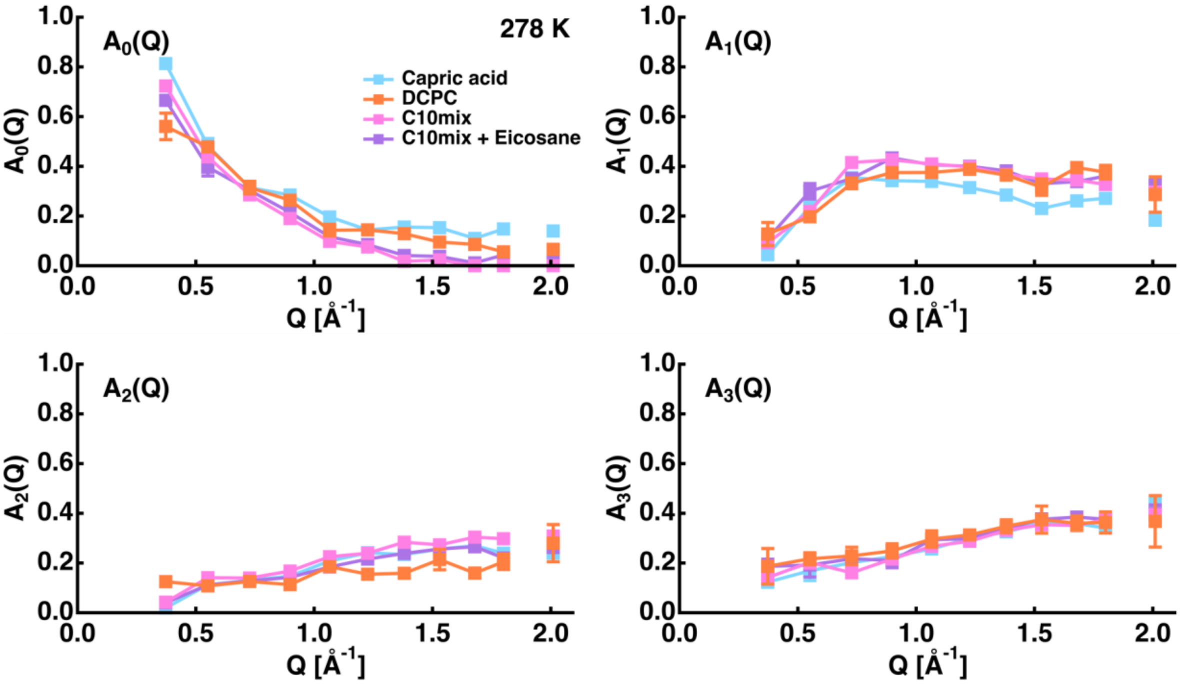


**Figure S1:** Amplitudes $A_0$ to $A_3$ of all samples at 278 K.

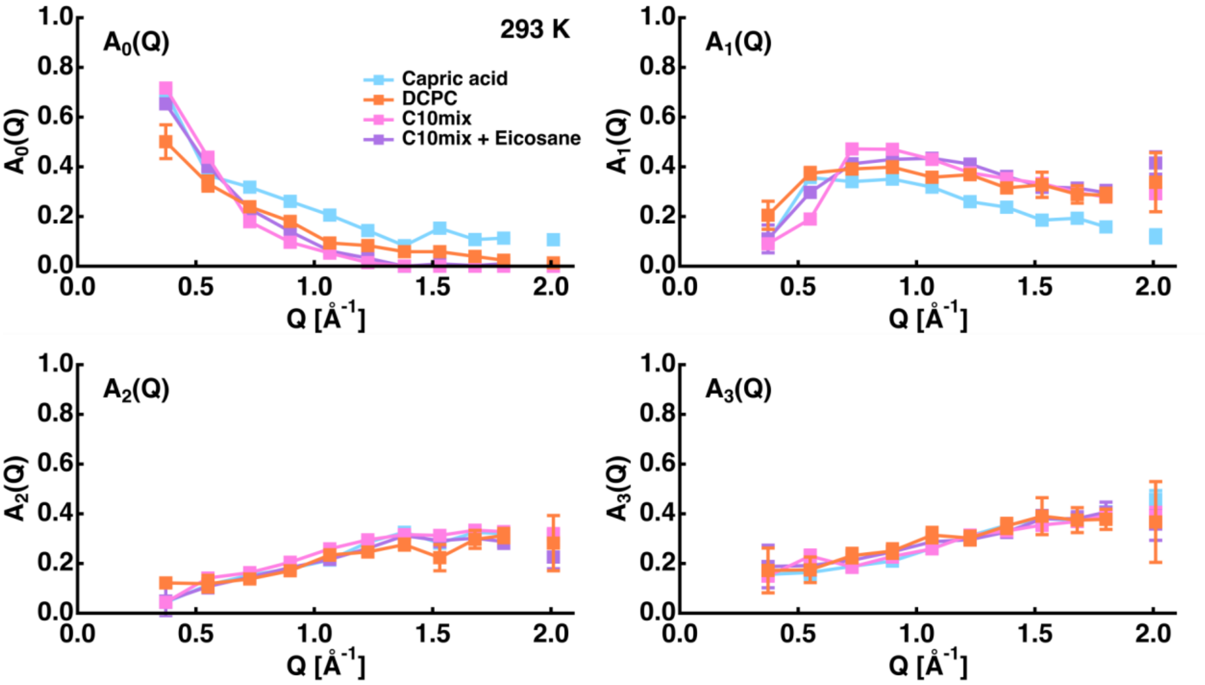


**Figure S2:** Amplitudes $A_0$ to $A_3$ of all samples at 293 K.

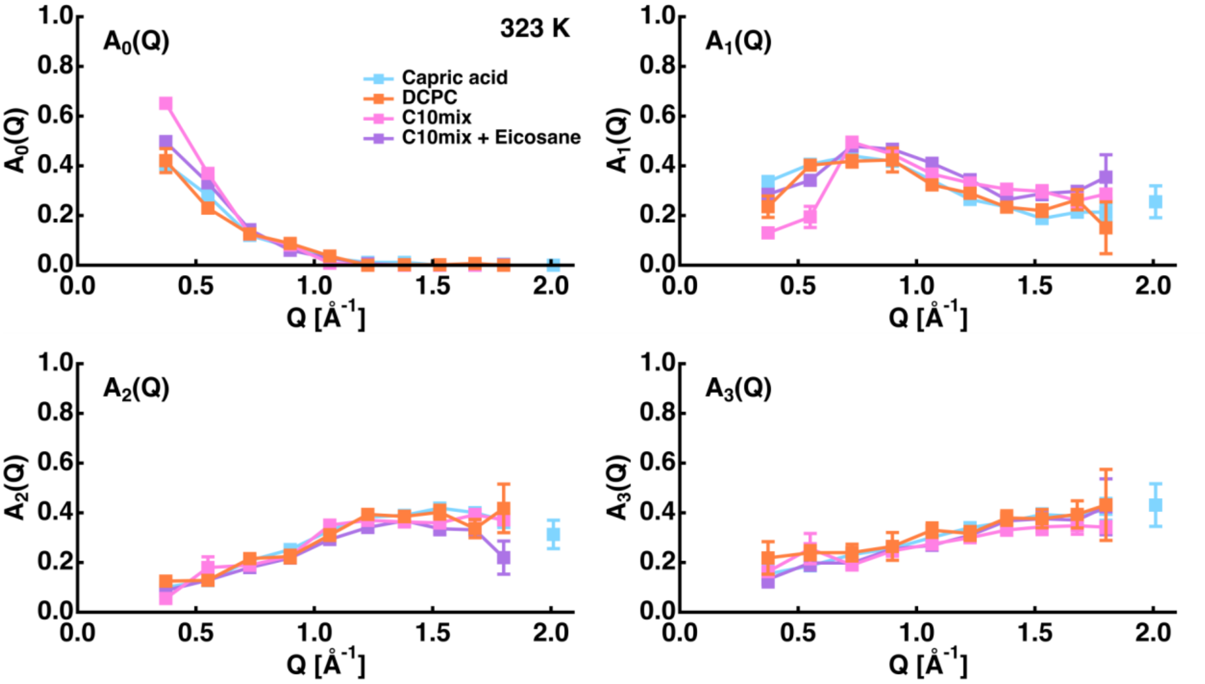


**Figure S3:** Amplitudes $A_0$ to $A_3$ of all samples at 323 K.

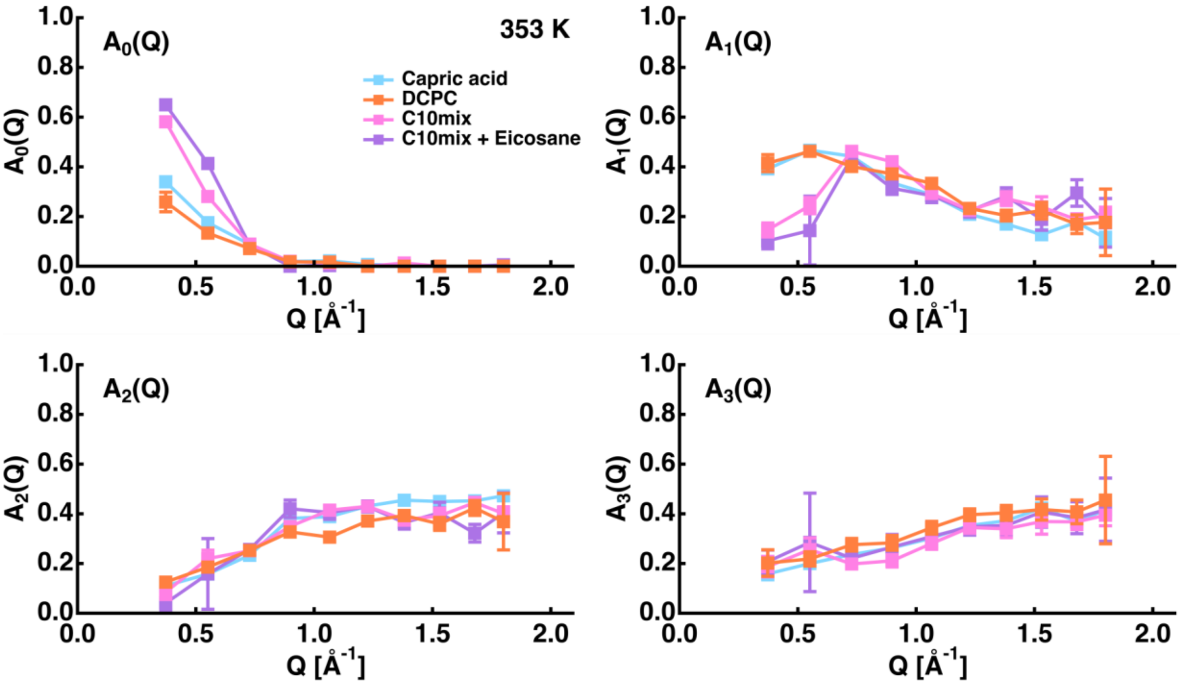


**Figure S4:** Amplitudes $A_0$ to $A_3$ of all samples at 353 K.

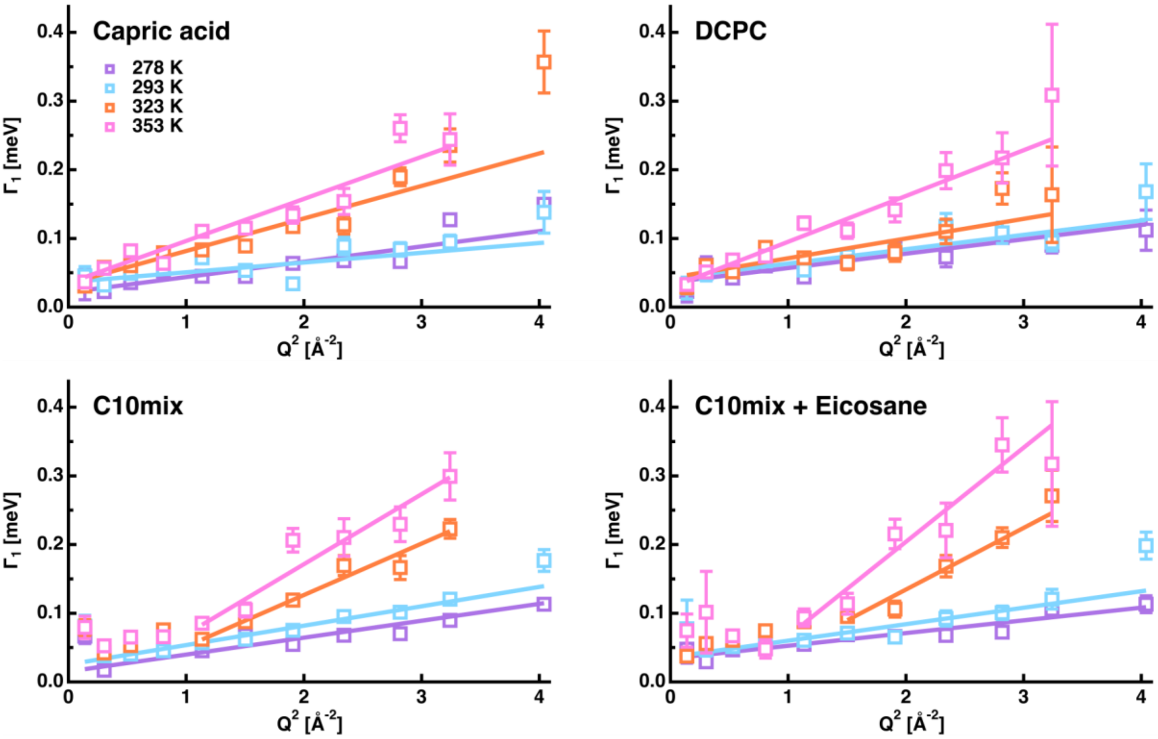

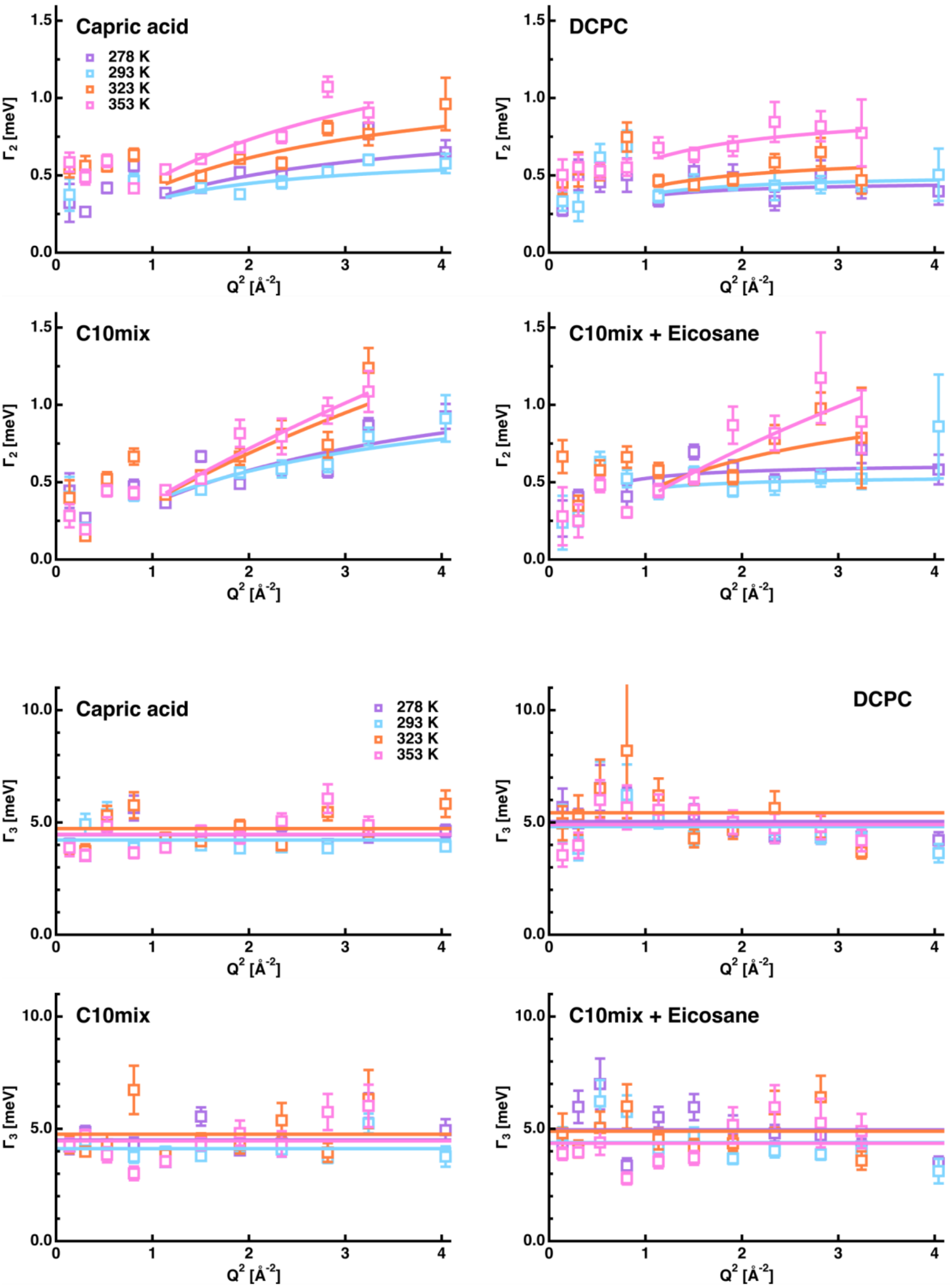


**Figure S5 – S7:** Fits of the HWHM $\Gamma_1$ to $\Gamma_3$ of all samples as function of $Q^2$.

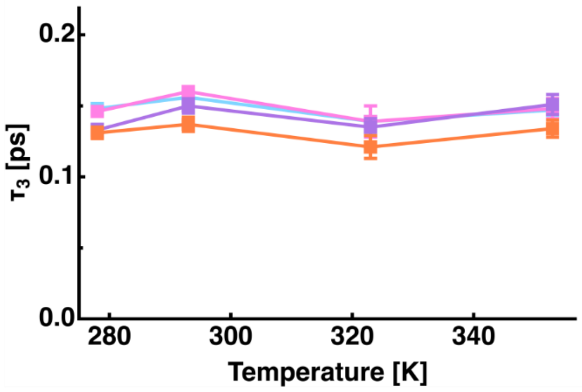


**Figure S8:** Rotational correlation time as a function of temperature.

Capric acid:

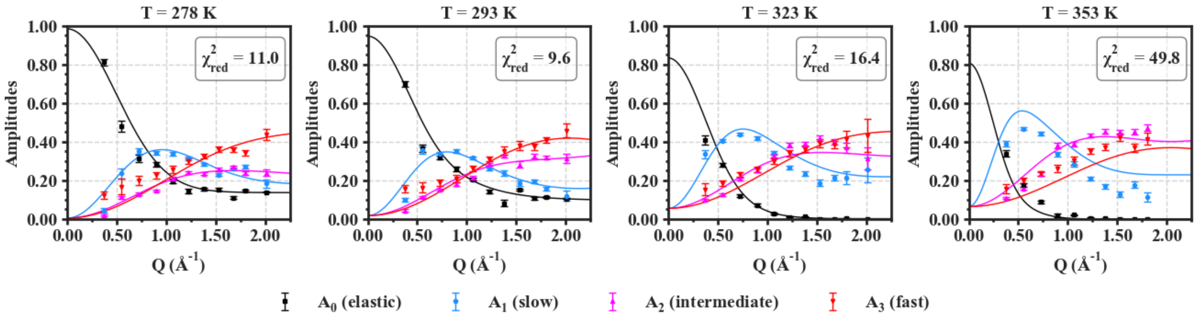


C10 mix:

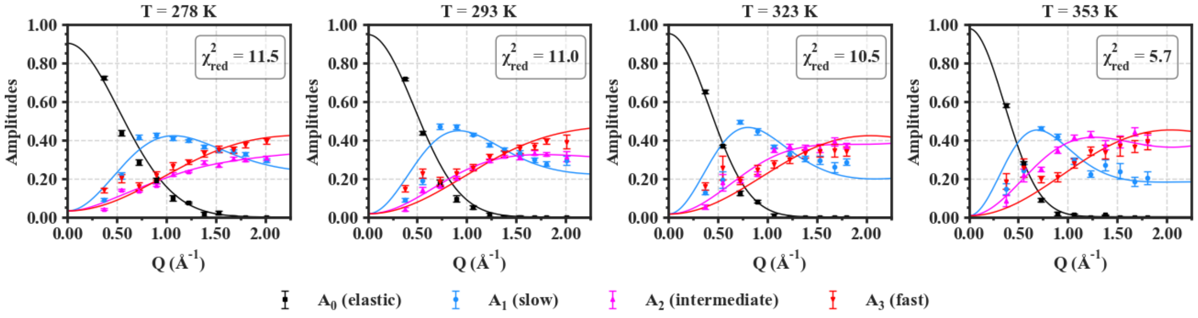


C10 mix + eicosane:

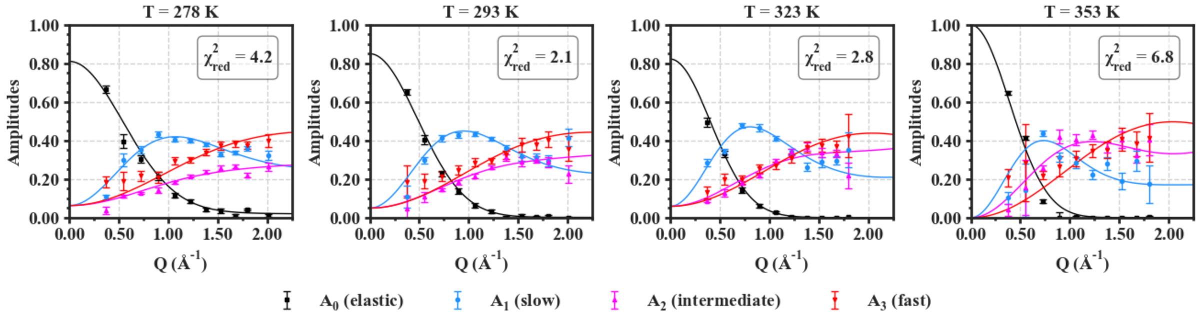


DCPC:

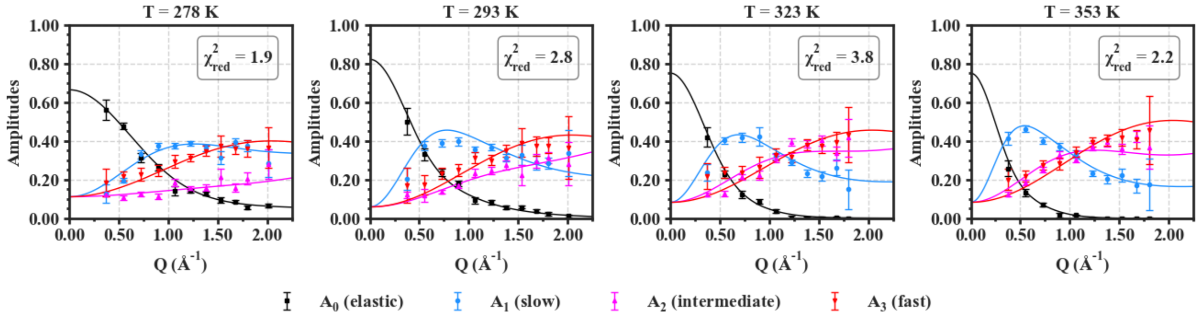


**Figure S9:** Fits of the amplitudes for all four samples. Error bars are within symbols if not shown.
